# Semantic Update as a Predictor of Reading Time: Moving Beyond Word-Level Surprisal

**DOI:** 10.64898/2026.10.12.758612

**Authors:** Siddharth Gupta, Alessandro Lopopolo, Milena Rabovsky

**Affiliations:** Department of Psychology, University of Potsdam, Karl-Liebknecht-Straße 24-25, 14476 Potsdam, Germany

**Keywords:** N400, sentence processing, semantic prediction, sentence gestalt model, eye tracking, reading times

## Abstract

This study explores the role of semantic prediction error in shaping reading behavior, as reflected in reading time during naturalistic reading. We operationalize semantic prediction error using semantic update (SU), a measure derived from the Sentence Gestalt (SG) model, which simulates incremental predictive comprehension at the level of sentence meaning. While SU has been previously linked to the N400 ERP component - an index of semantic processing - here we test its behavioral relevance by examining how it predicts total word-fixation durations in an eye-tracking corpus. To contextualize the role of semantic prediction error, we compare SU with lexical prediction error, quantified as next-word surprisal estimated from a language model (LM) trained on the same corpus as the SG model.

Our results show that SU significantly contributes to explaining total word-fixation durations, even after accounting for LM-based surprisal and lower-level lexical factors. We also find that this predictive relationship holds primarily for content words, but not for function words, which parallels findings for lexical surprisal.

These findings support the view that semantic prediction error plays a unique and measurable role in guiding real-time reading behavior, complementary to lexical prediction.

## 1 Introduction

Reading is a complex cognitive skill that unfolds through several partly interrelated processes, including prediction (Wong et al., 2024), word recognition (Sereno, 2003), lexical access (Segalowitz & Lane, 2000), syntactic parsing (Lopopolo et al., 2019), and semantic integration (Hagoort et al., 2009). Understanding these processes offers valuable insights into the cognitive mechanisms underlying language comprehension. By linking metrics derived from computational models to real-time human reading behavior, researchers can test mechanistic theories of language processing.

Advances in language modeling have fueled interest in using computational measures to explain reading behavior observed in eye-tracking studies. One key metric is surprisal, which reflects the unexpectedness of a word given its preceding context. Surprisal has been at the forefront of the debate as a primary metric for sentence processing difficulty, largely attributed to expectation-centered theories of comprehension (Hale, 2001; Levy, 2008). However, because language models are fundamentally optimized solely to just predict the next sequential lexical item during an input, surprisal does not explicitly track how a reader continuously updates their internal representation of the events described in the sentence.

To capture this meaning-level update, recent work has introduced Semantic Update (SU), a metric derived from the Sentence Gestalt (SG) model (McClelland et al., 1989). The SG neural network is specifically designed to map sequences of incoming words to a predictive representation of sentence meaning including the assignment of concepts to thematic roles in the described event(Rabovsky et al., 2018). Rather than merely tracking context-dependent lexical prediction errors, SU represents the amount of change in the model’s internal predictive representation of meaning resulting from encountering a new word. Given a specific context, the model develops expectations for the overall sentence meaning. If an incoming word is compatible with these expectations, the information gain is minimal, and no significant adjustment is needed to the model’s internal representation of meaning. Conversely, an unexpected word triggers greater information gain, requiring a higher degree of adjustment to accommodate the new semantic information. Unexpectedness of words in language is graded, and gradually larger unexpectedness will result in gradually larger adjustments to the model’s internal semantic representation. The theoretical validity of SU is supported by its success in modeling neurophysiological responses: it successfully predicts the N400 event-related potential (ERP) component (Rabovsky et al., 2018), the most widely studied ERP component in language comprehension research (Kutas & Federmeier, 2011). Notably, SU predicts N400 amplitudes beyond the variance explained by surprisal (Lopopolo & Rabovsky, 2024).

Given the relevance of N400 during semantic processing (Kutas & Federmeier, 2011), it is plausible that the semantic update processes captured by SU will also manifest through behavioral measures such as reading times. The link between N400 and human reading times is grounded in evidence that, though operating on different timelines, both are sensitive to the same linguistic constraints (Dambacher & Kliegl, 2007). Additional findings in the eye-tracking literature have demonstrated that semantic-based measures significantly impact reading times (Kun et al., 2023). Following the theoretical and neurophysiological links, this study utilizes SU to predict per-word total fixation duration during naturalistic reading.

Beyond general semantic update and prediction error, it is interesting to understand how these predictive effects are modulated by syntactic category - namely, content and function words. Content words are those that carry the core meaning of a sentence. They typically denote entities, actions, properties, or states – for example, nouns (dog, teacher), verbs (run, give), adjectives (blue, difficult), and adverbs (quickly, often). These words provide reference and denotation, anchoring linguistic expressions to concepts in the world and contributing directly to the propositional content of a sentence. In contrast, function words (e.g., articles, prepositions, conjunctions, auxiliary verbs) serve primarily grammatical and relational purposes. They signal how content words combine, mark syntactic roles, and structure the sentence without introducing substantial new concepts themselves. While indispensable for fluent comprehension, the contribution of function words is less about reference and more about scaffolding the semantic frame in which content words operate. Frank et al. (2013) demonstrated a distinct dissociation in N400 neural activity: surprisal reliably predicts the N400 for content words, whereas no such relation was found for function words. Given that SU effectively models the N400 (Rabovsky et al., 2018), it seems relevant to investigate whether the effects of SU on reading times exhibit a similar dissociation. Furthermore, recent computational research highlights that massive language models struggle to accurately model reading times across these categories, systematically under-predicting the time spent on content words due to over-memorization of training data (Oh & Schuler, 2022). This raises the question of whether models tracking Semantic Update exhibit similar biases.

### 1.1 Current Study

In the current study, we hypothesize that real-time reading behavior reflects both the lexical-level prediction of individual word forms (measured via surprisal) and the incremental prediction error at the level of sentence meaning (measured via SU). As each word is encountered, it not only possesses a context-dependent predictability at the lexical level but also modifies the evolving predictive representation of sentence meaning. Furthermore, we investigate whether the effect of SU varies across syntactic categories, potentially playing a more prominent role in the processing of content words than function words.

To test these hypotheses, we examined the effect of SU as a predictor of total reading time using an eyetracking dataset of 46 native English-speaking participants who read naturalistic texts in English (Siegelman et al., 2022). We fit a linear mixed-effects model which includes LM-based-surprisal and lexical features (i.e. word length, position and frequency), alongside SU, predicting per-word total reading time duration. Crucially, to ensure a rigorous comparison, the LM used to calculate surprisal was trained on the same dataset as the SU model. Lastly, to test whether predictive effects on reading time are modulated by a word’s class, we analyzed the interaction between word type and the computational metrics of interest. Overall, this study aims to deepen the understanding of how incremental semantic construction shapes real-time reading behavior. Finally, to understand how these models deviate from human behavior, we conduct a residual analysis. This allows us to test whether our models replicate the under-prediction biases for content words observed in the massive language models (Oh & Schuler, 2022), or if they reflect the cognitive load of meaning-bearing words differently.

## 2 Materials and Methods

### 2.1 Participants and Eye-tracking Dataset

This study utilized the Multilingual Eye-movement Corpus (MECO) (Siegelman et al., 2022), a large cross-linguistic eye-tracking corpus designed to support research on human sentence reading. The MECO corpus includes eye-tracking data collected from participants reading texts in 13 different languages. Each participant read 12 Wikipedia-style texts in their dominant language (L1), while their eye movements were recorded using an EyeLink 1000/1000+ eye-tracker (in case of the English subset) at a sampling rate of 1000 Hz. The dataset contains a variety of eye-tracking measures obtained during the reading task, such as reading time and regression indicators.

Pre-processing involved several steps to ensure data quality and comparability across measures. Extremely short total reading times (below 80 ms) along with durations greater than the 99th percentile (within each participant) were removed. The same procedure was applied to the number of fixations per word, with trials above the 99th percentile excluded. After filtering, the reading times were log-transformed, a standard practice in psycholinguistics studies. All the predictors were centered and scaled. For detailed information on participant recruitment, experimental setup, text selection, etc., refer to Siegelman et al. (2022).

For our analysis, we focused on the English subset of the dataset, which comprises data from 46 university student participants (mean age: 21.04). We further filtered the dataset to include only sentences composed of words present in the SG model’s training vocabulary, excluding any sentences containing acronyms. The resulting dataset included 53 sentences drawn from the 12 texts, consisting of 26,876 observations.

### 2.2 Computational Models and Predictors

In this study we adopt two distinct computational models – a language model (LM) and a Sentence Gestalt (SG) model – which optimize different predictive objectives. A standard LM predicts surface forms, with surprisal indexing the unexpectedness of the next token. By contrast, the SG model predicts structured event semantics. This functional distinction – token-level prediction vs. event-level meaning prediction – explains why SU should track sentence-level meaning, while surprisal captures lexical predictability across tokens. To ensure differences in performance are driven by training task rather than training data exposure, both models were trained on the identical 8-million-token Rollenwechsel-English (RW-eng) corpus (Sayeed et al., 2018). Figure 2 gives an overview of the modeling and analysis pipeline.

#### 2.2.1 The Sentence Gestalt (SG) Model

The SG model (McClelland et al., 1989) maps sentences to the description of their meaning, represented as a set of role-filler pairs, where roles (e.g., agent, patient, recipient) specify thematic functions in the described event, and fillers specify the entities fulfilling these roles. The model is probabilistic in nature; it approximates a function that computes a conditional probability distribution between the sequence of words that make up the sentence and the corresponding role-filler pairs; the model is also predictive. During training, after each successive word in a sentence is encountered, it is required to predict the complete set of role–filler pairs for the sentence, including fillers that have not yet been encountered in the input. Consequently, the model continuously updates its expectations about the event structure as each new word is processed. For example, in the sentence, “The teacher gave the student a book”, after processing the partial input “The teacher gave…”, the model infers that “teacher” fills the agent role and that “gave” corresponds to the action, while simultaneously generating expectations about probable recipients and patients based on the learned statistical regularities. These predictions are progressively refined as the remainder of the sentence unfolds.

The SG model comprises two components, namely, an update network (encoder) and a query network (decoder), see Figure 1a. The update network processes each incoming word sequentially to modify the internal representation of sentence meaning, referred to as the Sentence Gestalt layer. This consists of a recurrent long short-term memory (LSTM) layer (Hochreiter & Schmidhuber, 1997) which integrates the current input word with the previous SG representation (*sg*_*t−*1_) to produce the updated representation (*sg*_*t*_).

**Figure 1:**
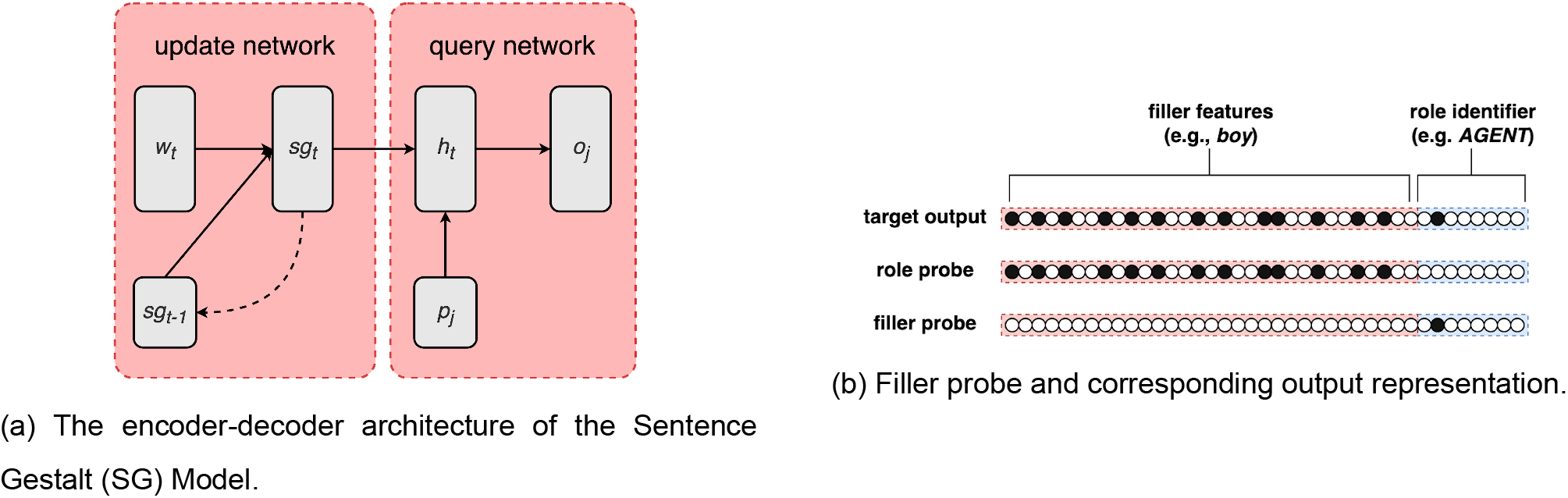
Role- and filler-based probing of the query network.

**Figure 2:**
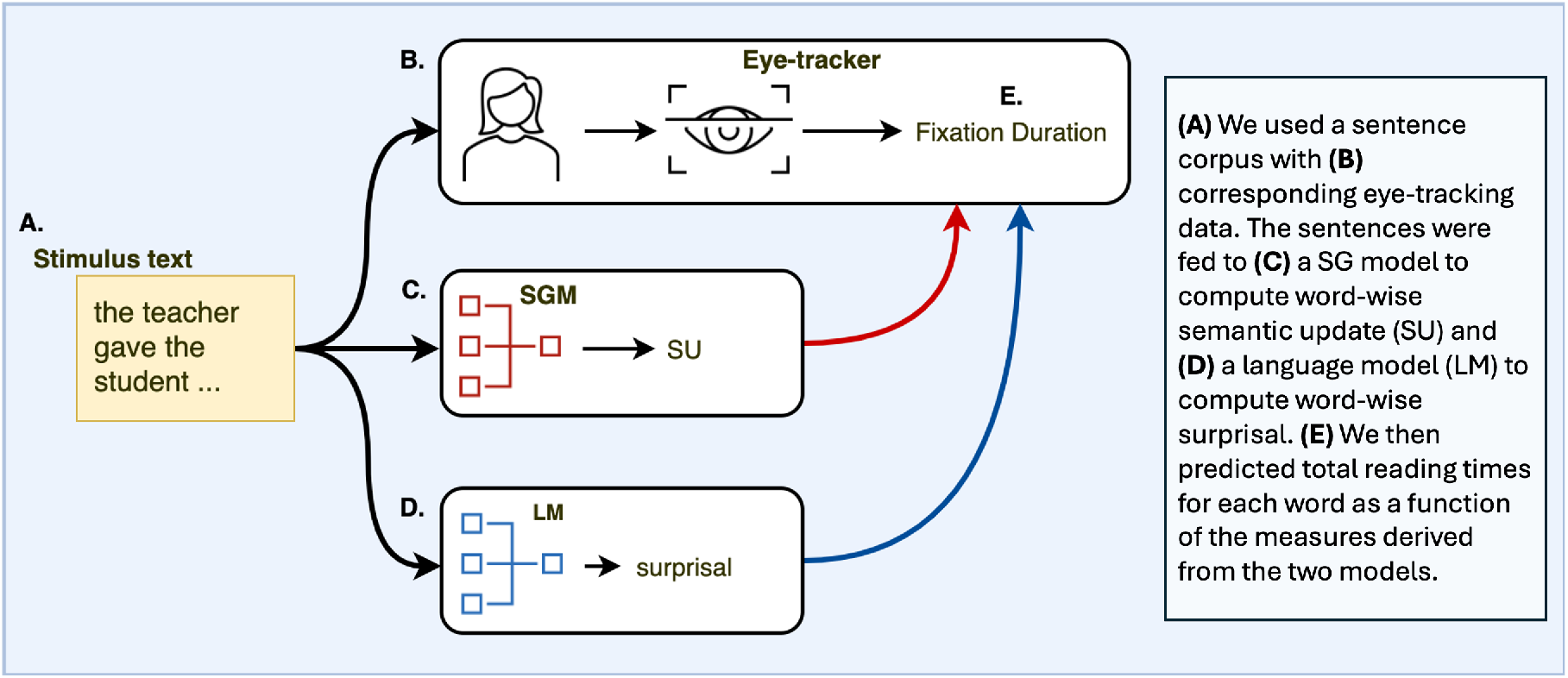
Visual abstract. Schematic overview of the experimental and modeling pipeline used to relate Sentence Gestalt (SG) semantic update and language-model surprisal to human reading times.

The query network retrieves event information from the SG representation and is primarily used during model training. For each role–filler pair in the event representation, the model receives a probe vector (*p*_*j*_) containing either a one-hot representation of the thematic role or the distributed semantic representation of the filler (Bojanowski et al., 2017). A hidden layer (*h*_*t*_) combines this probe with the output of the update network to generate an output vector (*o*_*j*_) representing the complete role–filler pair. Thus, when provided with a role probe, the model predicts the corresponding filler; when provided with a filler probe, it predicts the corresponding role. In both cases, the output contains the filler’s distributed semantic features together with one-hot representations of the thematic role and frame number. Figure 1b illustrates the representations of a role probe and a filler probe, together with the corresponding complete role–filler representation produced at the output layer.

Through this bidirectional probing procedure, the model learns probabilistic associations between event roles and their fillers.

#### 2.2.2 The Language Model

While the SG model focuses on sentence meaning (operationalized as role-filler pairs), LMs instead work at predicting individual tokens in a sentence as a function of their preceding context. The LM architecture is similar to the update network of the SG model: in both architectures, an LSTM recurrently integrates each incoming word with the representation of the preceding context. The main difference lies in how the LSTM representation is used. In the SG model, it constitutes the Sentence Gestalt representation and is passed to the query network, which retrieves role–filler information. In the LM, by contrast, the LSTM feeds into an output network that generates a probability distribution over the possible next words in the sequence.

For a given sequence of *n* words, ⟨*w*_1_, *w*_2_, *w*_3_, …, *w*_*n*_⟩, the task of a language model is to predict the most likely continuation *w*_*n*+1_ based on the statistical patterns learned from the corpus on which it was trained. For example, given the partial sequence “The teacher gave the student a”, the LM would be expected to assign a higher probability to lexical items such as “book” or “homework” as compared to “spoon” or “shoe”.

#### 2.2.3 Model training

Both the SG model and the LM were trained on the Rollenwechsel-English (RW-eng) corpus (Sayeed et al., 2018), comprising approximately 8 million tokens. The corpus provides sentence-level event annotations based on PropBank semantic roles (Palmer et al., 2005), representing events as predicates and role-filler pairs.

Crucially, the two models were trained on the same sentences but with different learning objectives. The SGM was trained to map word sequences onto their corresponding role-filler representations, whereas the LM was trained on next-word prediction using only the word sequences. Thus, differences between SU and lexical surprisal cannot be attributed to differences in the training data, but reflect the distinct learning objectives of the two models.

### 2.3 Computational predictors

#### 2.3.1 Semantic Update

When processing each word of a sentence, the SG model updates its sentence gestalt (SG) layer, which represents the model’s current expectation about sentence meaning. The magnitude of this update – how much the network’s internal state changes from one word to the next – can be directly quantified from the layer activation and is called semantic update (SU).

Formally, SU, as reflected by the presentation of a new word *w*_*t*_, is defined as follows:

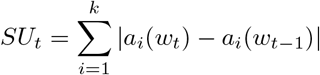

where *a*_*i*_(*w*_*t*_) is the activation at the *i*-th unit in the gestalt layer (with *k* units) as the network encounters the *n*-th word in the sentence. Hence, SU captures the total change in activation by summing the absolute differences across SG layer units before and after encountering a word.

#### 2.3.2 Lexical surprisal

Surprisal measures the level of unexpectedness of a word based on the context provided by the preceding words in the sentence. It is defined as the negative logarithm of the probability of the word given its context: *S*(*w*_*t*+1_|*w*_1_, *w*_2_, …, *w*_*t*_) = − log *P* (*w*_*t*+1_|*w*_1_, *w*_2_, …, *w*_*t*_), where *S*(*w*_*t*+1_|*w*_1_, *w*_2_, …, *w*_*t*_) represents the surprisal of the word *w*_*t*+1_ given the preceding sequence of words *w*_1_, *w*_2_, …, *w*_*t*_, and *P* (*w*_*t*+1_|*w*_1_, *w*_2_, …, *w*_*t*_) is the conditional probability of *w*_*t*+1_ given the sequence *w*_1_, *w*_2_, …, *w*_*t*_.

### 2.4 Analyses

#### 2.4.1 Total fixation duration

We fit a linear mixed-effects model to predict log-transformed per-word total reading duration as a function of semantic update, LM-based-surprisal, and low-level features, namely, word length, word position within the sentence, and word frequency (Norvig, 2008; Tatman, 2017). To account for variability across participants and items, we included random intercepts and random slopes for semantic update and LM-based-surprisal by participant and by word. According to our hypothesis, larger update in the SG layer, operationalised by SU, should result in longer reading durations.

#### 2.4.2 Word type-based analysis

We also ran mixed-effects models similar to 2.4.1 on two subsets of the data, based on syntactic word category: content words and function words. For this, we divided the dataset into content words (19,181 observations) and function words (7,695 observations). Content words were defined as adjectives, nouns, verbs, and adverbs, based on part-of-speech (POS) tags following the annotation scheme of Marcus (1993). We expect SU to be a better predictor of total reading time for content words than function words, due to the model’s focus on tracking semantic prediction error (Lopopolo & Rabovsky, 2024).

To assess the robustness and significance of the difference in SU’s prediction performance across the two word categories, we ran a mixed-effects model including word type (content vs. function) as an interaction term, along with its interactions with LM-based surprisal and SU.

We then evaluated how well SU and LM-based surprisal predict word duration for each word category by examining their residuals. We fit two separate models: one including the predictor SU, and one including LM-based surprisal, both with the same control variables and random effects as in 2.4.1. Residuals were computed as the difference between the observed durations and the model predictions. An over-prediction would indicate that the model expected a word to cause more processing difficulty, and thus take longer to read, than it actually did for human readers. Conversely, an under-prediction would indicate that the model underestimated the human reading time. This builds on the findings from (Oh & Schuler, 2022), that demonstrated that massive language models systematically under-predict reading times for content words due to their over-memorization of training data, while over-predicting for function words to mathematically compensate. By analyzing the directionality of the residuals across syntactic categories, we can determine whether our computationally distinct models exhibit the same biases or if they capture the cognitive load of the meaning-bearing words differently. Subsequently, we ran separate t-tests for each model and each word category on the respective residuals to test whether residual values are significantly different from zero. Lastly, to assess whether there is a significant difference between the mean residuals of content words and function words within the same model, we ran Welch’s t-tests.

## 3 Analyses and Results

### 3.1 Total fixation duration

The results in Table 1 indicate that SU significantly predicts total per-word fixation duration (*β* = 0.03, *t* = 2.81, *p <* 0.001) beyond the influence of LM-based surprisal. This indicates that larger updates of the SG layer representation correspond with longer reading times for the respective word even when controlled for other relevant factors.

**Table 1:**
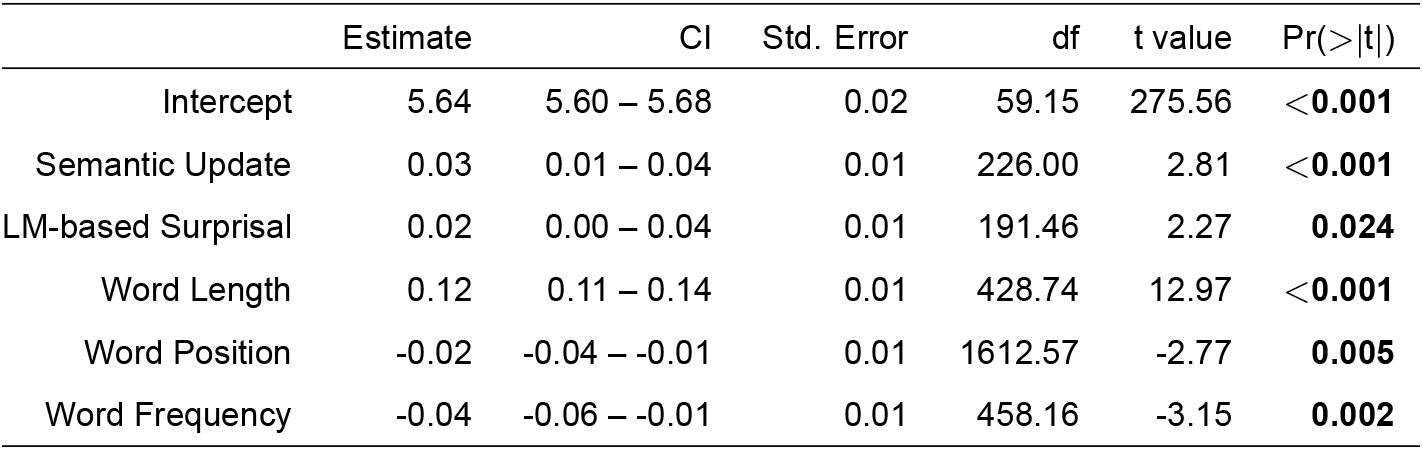
Results of the linear mixed-effect model predicting total per-word fixation duration as a function of semantic update, LM-based surprisal, and word-level features.

|  | Estimate | CI | Std. Error | df | t value | Pr(>\|t\|) |
| --- | --- | --- | --- | --- | --- | --- |
| Intercept | 5.64 | 5.60 – 5.68 | 0.02 | 59.15 | 275.56 | <0.001 |
| Semantic Update | 0.03 | 0.01 – 0.04 | 0.01 | 226.00 | 2.81 | <0.001 |
| LM-based Surprisal | 0.02 | 0.00 – 0.04 | 0.01 | 191.46 | 2.27 | 0.024 |
| Word Length | 0.12 | 0.11 – 0.14 | 0.01 | 428.74 | 12.97 | <0.001 |
| Word Position | -0.02 | -0.04 – -0.01 | 0.01 | 1612.57 | -2.77 | 0.005 |
| Word Frequency | -0.04 | -0.06 – -0.01 | 0.01 | 458.16 | -3.15 | 0.002 |

As expected, LM-based surprisal also has a significant impact in predicting the reading times (*β* = 0.02, *t* = 2.27, *p <* 0.024). All low-level word features, word length (*β* = 0.12, *t* = 12.97, *p <* 0.001), word position (*β* = −0.02, *t* = −2.77, *p* = 0.005) and word frequency (*β* = −0.04, *t* = −3.15, *p* = 0.002) as well have a significant relationship with the total per-word fixation durations, in the expected directions, i.e., longer and more surprising words increase total fixation durations, whereas highly frequent words and words appearing later in the sentence decrease them.

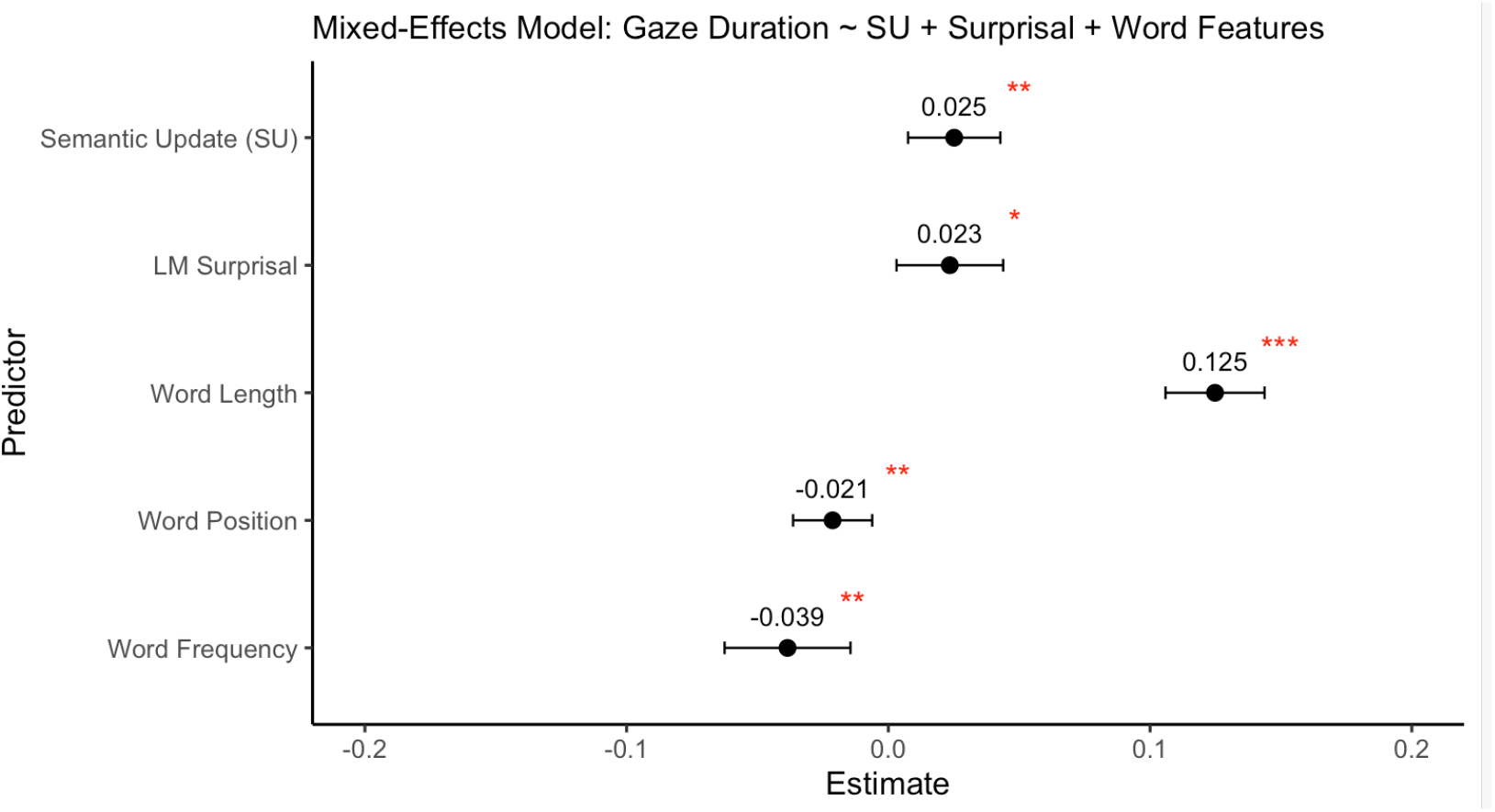

### 3.2 Subset Analysis based on word type

#### 3.2.1 Content words

The results for the content-word subset analysis with LM-based surprisal and Semantic Update (SU) are presented in Table 2. SU showed a significant association with the total per-word fixation duration (*β* = 0.03, *t* = 2.41, *p* = 0.017). LM-based surprisal also had a strong effect (*β* = 0.05, *t* = 3.52, *p <* 0.001), and word length showed a substantial effect (*β* = 0.12, *t* = 11.84, *p <* 0.001), along with word position (*β* = −0.04, *t* = −3.49, *p* = 0.001). On the other hand, word frequency (*β* = −0.03, *t* = −1.13, *p* = 0.26) showed no significant association.

**Table 2:**
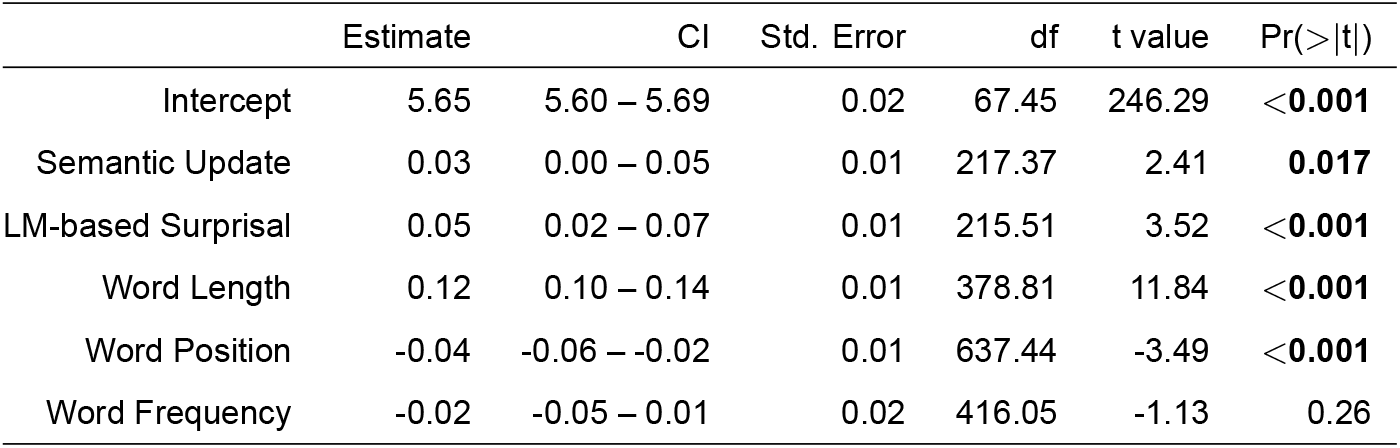
Results of the linear mixed-effect model predicting total per-word fixation duration as a function of semantic update (SU), LM-based surprisal, and word-level features on dataset subsetted for content words.

#### 3.2.2 Function words

Results for the function-word subset analysis are presented in Table 3. For function words, total per-word fixation durations were primarily predicted by word length (*β* = 0.11, *t* = 3.21, *p* = 0.002). Semantic Update (*β* = 0.00, *t* = 0.18, *p* = 0.85), LM-based surprisal (*β* = −0.01, *t* = −0.94, *p* = 0.35), word position (*β* = 0.00, *t* = 0.4, *p* = 0.69) and word frequency (*β* = −0.05, *t* = −1.49, *p* = 0.15) did not have a significant effect.

**Table 3:** Results of the linear mixed-effect model predicting total per-word fixation durations as a function of semantic update (SU), LM-based surprisal, and word-level features on dataset subsetted for function words.

|  | Estimate | CI | Std. Error | df | t value | Pr(>\|t\|) |
| --- | --- | --- | --- | --- | --- | --- |
| Intercept | 5.60 | 5.55 – 5.66 | 0.03 | 107.30 | 202.67 | <0.001 |
| Semantic Update | 0.00 | -0.03 – 0.03 | 0.01 | 29.85 | 0.18 | 0.85 |
| LM-based Surprisal | -0.01 | -0.04 – 0.01 | 0.01 | 50.38 | -0.94 | 0.35 |
| Word Length | 0.11 | 0.04 – 0.17 | 0.03 | 48.22 | 3.21 | 0.002 |
| Word Position | 0.00 | -0.02 – 0.03 | 0.01 | 1386.81 | 0.40 | 0.69 |
| Word Frequency | -0.05 | -0.11 – 0.01 | 0.03 | 38.13 | -1.49 | 0.15 |

#### 3.2.3 Interaction model

The interaction coefficients (Table 4) show that the effect of SU on total per-word fixation duration is smaller for function words than for content words (*β* = −0.05, *t* = −2.52, *p* = .013). Likewise, the effect of LM-based surprisal is also smaller for function words (*β* = −0.03, *t* = −2.02, *p* = .046) in comparison. The negative coefficients indicate that these predictors have stronger effects for meaning-bearing content words.

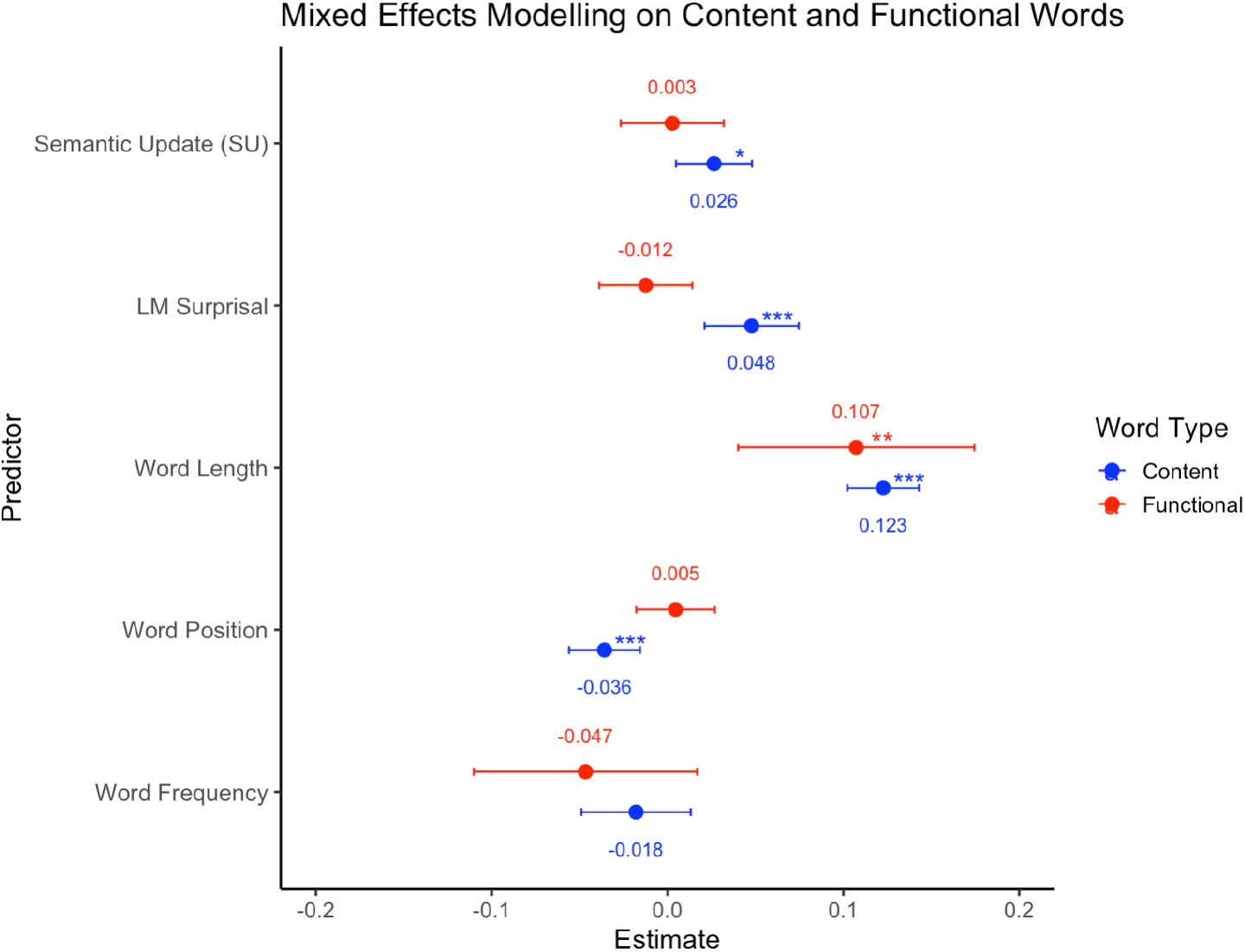

**Table 4:** Results of the linear mixed-effect model predicting total per-word fixation duration as a function of semantic update, LM-based surprisal, word-level features and interaction term for word type with reference level set to content words.

|  | Estimate | CI | Std. Error | df | t value | Pr(>\|t\|) |
| --- | --- | --- | --- | --- | --- | --- |
| Intercept | 5.65 | 5.61 – 5.69 | 0.02 | 68.10 | 266.13 | <**0.001** |
| Semantic Update | 0.04 | 0.02 – 0.05 | 0.01 | 257.15 | 3.63 | <**0.001** |
| LM-based Surprisal | 0.04 | 0.01 – 0.06 | 0.01 | 357.83 | 2.94 | **0.004** |
| Word Length | 0.12 | 0.10 – 0.14 | 0.01 | 431.70 | 12.61 | <**0.001** |
| Word Position | -0.02 | -0.04 – -0.01 | 0.01 | 1626.01 | -2.62 | **0.009** |
| Word Frequency | -0.02 | -0.05 – 0.00 | 0.01 | 511.86 | -1.64 | 0.1 |
| Word Category | -0.06 | -0.10 – -0.01 | 0.02 | 601.67 | -2.26 | **0.024** |
| Semantic Update:Word Category | -0.05 | -0.09 – -0.01 | 0.02 | 169.15 | -2.52 | **0.013** |
| LM-based Surprisal:Word Category | -0.03 | -0.06 – -0.00 | 0.02 | 85.22 | -2.02 | **0.046** |

### 3.3 Residual Analysis

The results of the residual analysis reported in Table 5(Semantic Update: *t* = −2.79, *p* = 0.005, LM-based Surprisal: *t* = −2.48, *p* = 0.013) indicate that both models significantly over-predict total per-word fixation durations for content words. Conversely, for function words, the results (Semantic Update: *t* = −0.37, *p* = 0.71, LM-based Surprisal: *t* = −0.57, *p* = 0.57) suggest no systematic deviation in residuals, though the difference between the residuals of the word categories does not significantly differ (Table 6).

**Table 5:** Residuals (observed minus predicted log durations) for content and function words tested against zero.

| Model | group | t_statistic | df | p_value |
| --- | --- | --- | --- | --- |
| Semantic Update | Content Words | -2.79 | 19180.00 | **0.005** |
| Semantic Update | Function Words | -0.37 | 7694.00 | 0.71 |
| LM-based Surprisal | Content Words | -2.48 | 19180.00 | **0.013** |
| LM-based Surprisal | Function Words | -0.57 | 7694.00 | 0.57 |

**Table 6:** Difference of residuals between content and function words tested against zero.

| Model | t_statistic | df | p_value |
| --- | --- | --- | --- |
| Semantic Update | -1.28 | 15360.10 | 0.2 |
| LM-based Surprisal | -0.94 | 15329.99 | 0.35 |

## 4 Discussion and Conclusions

The present study investigated whether Semantic Update (SU) – measure of sentence-level semantic prediction error derived from the Sentence Gestalt model (Lopopolo & Rabovsky, 2024; Rabovsky et al., 2018) – explains variation in natural reading times beyond lexical surprisal (Levy, 2008). We found that SU predicted total fixation duration and explained unique variance after controlling for surprisal and standard lexical variables. Notably, these effects were stronger for content words compared to function words.

SU’s contribution to reading times beyond lexical surprisal suggests that models based solely on next-word prediction do not fully account for the observed variation in natural reading behavior. Instead, readers appear to be sensitive not only to the predictability of the upcoming lexical items but also to the changes in the evolving predictive representation of sentence meaning. In light of expectation-based theories of comprehension (Hale, 2001; Levy, 2008), our findings suggest that predictive processing operates at multiple representational levels, in line with the idea of hierarchical prediction and prediction error processing in the brain (Friston, 2005). While lexical surprisal effectively captures expectations regarding upcoming word forms, SU indexes prediction error over the broader sentence meaning, highlighting a multidimensional updating process during reading. Furthermore, our findings are consistent with previous demonstration that SU predicts N400 amplitude beyond lexical surprisal (Lopopolo & Rabovsky, 2024; Rabovsky et al., 2018). Together, these results suggest that the computational processes captured by SU may underlie both neural and behavioral measures of language comprehension.

The selective effect of SU on content words suggests that semantic prediction might be more closely tied to conceptual processing rather than general processing difficulty during reading. This dissociation may arise because content words introduce the core concepts that drive sentence-level event representations. However, it should be noted that this dissociation might be partly an artifact of the SG model’s training procedure. Because the model is trained probing exclusively for semantic role-filler pairs, it is not explicitly required to generate and update predictions based on syntactic function words which don’t usually act as semantic fillers and are therefore not probed, but might nonetheless sometimes contribute to updating semantic expectations. At the same time, it is interesting to not that lexical surprisal exhibited a similar dissociation. The processing of function words was instead shaped solely by word length. The residual analysis strengthens this distinction and in addition offers a contrast to previous findings with massive language models. While (Oh & Schuler, 2022) found that large models systematically under-predict reading times for content words, potentially due to data memorization, both SU and lexical surprisal in our models over-predict them. This indicates our models are highly sensitive to the content words, predicting more processing time for them than humans actually require. Finally, a limitation of the current study is that the analysis was restricted to an English corpus; it remains to be further explored whether the effect of SU generalizes to languages with different morphological and syntactic typologies. Nevertheless, the present findings provide first evidence that sentence-level semantic prediction and updating contribute to natural reading behavior beyond lexical predictability. By integrating computational modeling with eye-tracking, the study supports a view of language comprehension in which prediction operates not only over individual words but also over an evolving predictive representation of sentence meaning.

## 5 Data Availability

The relevant data and analyses code will be made available upon publication to preserve anonymity during the review process.

